# Genotypic clustering does not imply recent tuberculosis transmission in a high prevalence setting: A genomic epidemiology study in Lima, Peru

**DOI:** 10.1101/418202

**Authors:** Avika Dixit, Luca Freschi, Roger Vargas, Roger Calderon, James Sacchettini, Francis Drobniewski, Jerome T. Galea, Carmen Contreras, Rosa Yataco, Zibiao Zhang, Leonid Lecca, Sergios-Orestis Kolokotronis, Barun Mathema, Maha R. Farhat

**Affiliations:** Boston Children’s Hospital, Boston MA; Harvard Medical School, Boston MA; Socios En Salud, Lima, Peru; Texas A & M University, College Station, TX; Imperial College, London, UK; University of South Florida, Tampa FL; Brigham and Women’s Hospital, Boston MA; SUNY Downstate School of Public Health, Brooklyn NY; Mailman School of Public Health, Columbia University, New York, NY; Massachussetts General Hospital, Boston, MA

## Abstract

**Background:** Whole genome sequencing (WGS) can elucidate *Mycobacterium tuberculosis* (Mtb) transmission patterns but more data is needed to guide its use in high-burden settings. In a household-based transmissibility study of 4,000 TB patients in Lima, Peru, we identified a large MIRU-VNTR Mtb cluster with a range of resistance phenotypes and studied host and bacterial factors contributing to its spread.

**Methods:** WGS was performed on 61 of 148 isolates in the cluster. We compared transmission link inference using epidemiological or genomic data with and without the inclusion of controversial variants, and estimated the dates of emergence of the cluster and antimicrobial drug resistance acquisition events by generating a time-calibrated phylogeny. We validated our findings in genomic data from an outbreak of 325 TB cases in London. Using a larger set of 12,032 public Mtb genomes, we determined bacterial factors characterizing this cluster and under positive selection in other Mtb lineages.

**Findings:** Four isolates were distantly related and the remaining 57 isolates diverged ca. 1968 (95% HPD: 1945-1985). Isoniazid resistance arose once, whereas rifampicin resistance emerged subsequently at least three times. Amplification of other drug resistance occurred as recently as within the last year of sampling. High quality PE/PPE variants and indels added information for transmission inference. We identified five cluster-defining SNPs, including *esxV* S23L to be potentially contributing to transmissibility.

**Interpretation:** Clusters defined by MIRU-VNTR typing, could be circulating for decades in a high-burden setting. WGS allows for an improved understanding of transmission, as well as bacterial resistance and fitness factors.

**Funding:** The study was funded by the National Institutes of Health (Peru Epi study U19-AI076217 and K01-ES026835 to MRF). The funding sources had no role in any aspect of the study, manuscript or decision to submit it for publication.

**Research in context:** *Evidence before this study:* Use of whole genome sequencing (WGS) to study tuberculosis (TB) transmission has proven to have higher resolution that traditional typing methods in low-burden settings. The implications of its use in high-burden settings are not well understood.

*Added value of this study:* Using WGS, we found that TB clusters defined by traditional typing methods may be circulating for several decades. Genomic regions typically excluded from WGS analysis contain large amount of genetic variation that may affect interpretation of transmission events. We also identified five bacterial mutations that may contribute to transmission fitness.

*Implications of all the available evidence:* Added value of WGS for understanding TB transmission may be even higher in high-burden vs. low-burden settings. Methods integrating variants found in polymorphic sites and insertions and deletions are likely to have higher resolution. Several host and bacterial factors may be responsible for higher transmissibility that can be targets of intervention to interrupt TB transmission in communities.

## Introduction

Tuberculosis (TB) remains among the top ten causes of deaths globally, with 10.4 million new cases in 2016 alone.^1^ Peru remains a high burden country for multidrug (MDR) TB with 117 TB cases were reported per 100,000 population in 2016 with approximately 9% being MDR or rifampicin resistant (RR).^1^ Molecular methods have been instrumental in tracing outbreaks and single nucleotide polymorphism (SNPs) identified using whole genome sequencing (WGS) have higher resolution in identifying transmission links when compared with traditional genotyping methods like spoligotyping or *Mycobacterium* interspersed repetitive unit-variable number tandem repeats (MIRU-VNTR).^2–12^ Yet we don’t yet fully understand how to use all the genetic information generated by WGS, by convention as much as 10% of the genome is excluded^2,3^ and in some instances too little remaining variation is found to aid resolution of transmission chains.^13^ Resolving transmission events accurately is particularly challenging in high-burden settings where multiple source case suspects are common. In addition to guiding public health interventions including appropriate contact tracing, identifying the source case can inform patient care in some cases such as in paediatric TB when the source case microbiological data can inform treatment.^14,15^ Further, in high-burden countries the term ‘outbreak’ may not apply as TB has been circulating continuously for decades or centuries.^16^ Given the renewed emphasis on active case finding^17^ and widespread adoption of sequencing, a closer look at use of WGS in high-burden settings is needed.

Control efforts against TB have been undermined by the emergence and spread of drug resistant TB (DR-TB). Current evidence suggests that most cases of DR TB are a result of transmission rather than de-novo evolution of the bacteria during treatment.^18–20^ Factors known to contribute to DR-TB transmission include delays in diagnosis and treatment,^21^ host factors like age, and immune status,^22–24^ as well as bacterial factors such as fitness and immunogenicity characteristics.^25–27^ It is well recognized that *Mycobacterium tuberculosis* (MTB) strains with the same DR conferring mutations have a range of fitness.^28–30^ However, to date few molecular fitness determinants have been characterized and seldom in the context of high transmissibility.^31,32^ Improved knowledge of such bacterial factors can inform efforts for transmission interruption by identifying targets for diagnosis, surveillance, and even potential therapeutics targeting fitness mechanisms. Here we use WGS data of the largest TB MIRU-VNTR cluster spanning pan-susceptible to MDR-TB isolates that was identified in 4,000 TB patients enrolled in a household transmissibility study. We examine both host and TB genotypic data to understand the evolution of isolates within this cluster, infer the timing of emergence of antibiotic resistance, and identify genetic bacterial factors unique to this cluster that may have contributed to its success.

## Methods

### Study Design

A TB household transmissibility and treatment outcome study was performed in northern Lima, Peru from September 2009 to August 2012. The study procedures including patient enrollment and consent have been previously described.^33,34^ Briefly, patients were enrolled if they were diagnosed with pulmonary TB (PTB) at public health clinics and were followed through therapy. Their household contacts were also followed with tuberculin skin testing and monitored for development of TB for a period of 12 months. The following were collected at time of TB diagnosis: clinical signs and symptoms, sociodemographic characteristics (e.g. age, gender, occupation, household type), geographical coordinates of household and health center, co-morbidities (HIV status, diabetes mellitus, renal disorder), as well as alcohol, tobacco and drug use.

Approval was obtained from the Research Ethics Committee of the Peruvian National Institute of Health (Lima, Peru) and the Committee on Human Studies at Harvard Medical School (Boston, MA).

Please refer to the supplement for bacterial culture, drug susceptibility testing (DST), DNA sequencing methodology and phylogenetic analysis.

## Results

### Patient and isolate characteristics

A large cluster of 148 strains with identical MIRU-VNTR pattern was identified. Majority of the patients in this cluster were male, HIV-negative and had multidrug resistant (MDR) TB (Table 1). About one-half were smear-positive (51.4%) and one-third used alcohol (35.8%) or other intoxicating substances (27.9%). Two patients had isolates that did not meet our sequencing quality criteria and were excluded. Of those patients with high quality sequence data (n=61) a higher majority were male with MDR-TB but were otherwise comparable to the superset of 148 (Table 1). Sequencing data revealed that 58 isolates belonged to the Latin America-Mediterranean LAM-4.3.3 sublineage, and three isolates were more distant and belonged to the sublineages X −4.1.1, T-4.8, and LAM-4.3.2. In the LAM-4.3.3 group we found 371 SNPs and 81 indels in total, of these 24 substitutions and one indel occurred in DR regions, and 42 substitutions and 23 indels in PE/PPE genes. With the exclusion of DR and PE/PPE regions, the average pairwise SNP difference between isolates was 21.69 (range: 0-84) and 22.7 (range:1-100), excluding and including the PE/PPE regions, respectively.

**Table 1:** Patient characteristics. None of the variables were significantly different between the patient with and without sequencing data using a t-test or a Chi-squared test.

|  | Cluster* | With WGS** |
| --- | --- | --- |
| Age*** | 32.06 (17-86) | 34.8 (17-86) |
| Female | 47 (31.8%) | 17 (27.9%) |
| Positive sputum smear | 76 (51.4%) | 30 (49.1%) |
| Heavy drinker in previous year | 53 (35.8%) | 24 (39.3%) |
| Drug use | 41 (27.9%)<br>(Total n = 147) | 20 (33.3%)<br>(Total n = 60) |
| HIV positive | 3 (2.1%)<br>(Total n = 140) | 3 (5.3%)<br>(Total n = 57) |
| Resistance Pattern<sup>#</sup> | N (%) | N (%) |
| Susceptible | 26 (17.6) | 9 (14.8) |
| Isoniazid Resistant | 18 (12.2) | 13 (21.3) |
| Multi-drug Resistant | 56 (37.8) | 31 (50.8) |
| Other | 36 (24.3) | 8 (13.1) |
| Not available | 12 (8.1) | 0 (0) |

\*N = 148 unless specified
\*\*N = 61 unless specified
\*\*\*Mean and range
<sup>#</sup>Resistance pattern for three of the sequenced strains was inferred based on mutations

The geographic distribution of strains based on household coordinates, colored by resistance pattern is shown in Figure 1. Comparison between genetic and geographic distance did not support that the cluster spread in a single geographic direction, even when three most distant strains were excluded (P = 0.2, Supplementary Figure 1).

**Figure 1:**
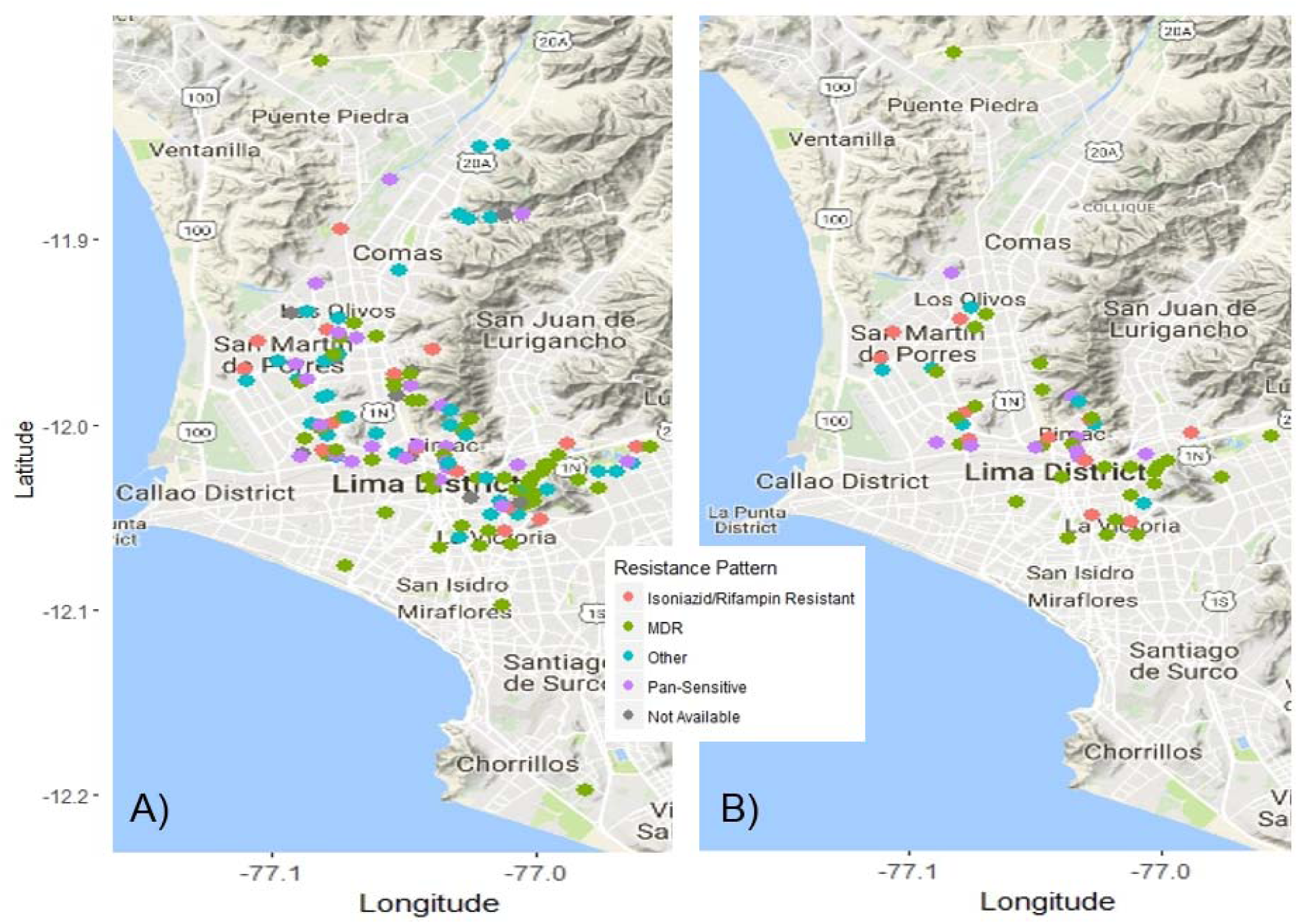
Geographic location of strains with resistance pattern. A) All 148 strains with identical MIRU-VNTR patterns, B) 61 strains that were sequenced.

The SNP-based phylogenetic tree (Figure 2, Supplementary Figure 2) demonstrated that the most genetically homogeneous group consisted of 57 isolates. These formed two main clades, where the first contained isolates that were pan-susceptible or streptomycin mono-resistant, and the second consisted of INH mono-resistant or MDR isolates. The origin of the 61 isolates was estimated around the middle of the 14^th^ century (1336 CE; 95% HPD 855-1680). The LAM-4.3 cluster of 58 isolates diverged ca 1923 (95% HPD 1856-1967) and innermost cluster of 57 isolates diverged ca 1968 (95% HPD 1945-1985).

**Figure 2:**
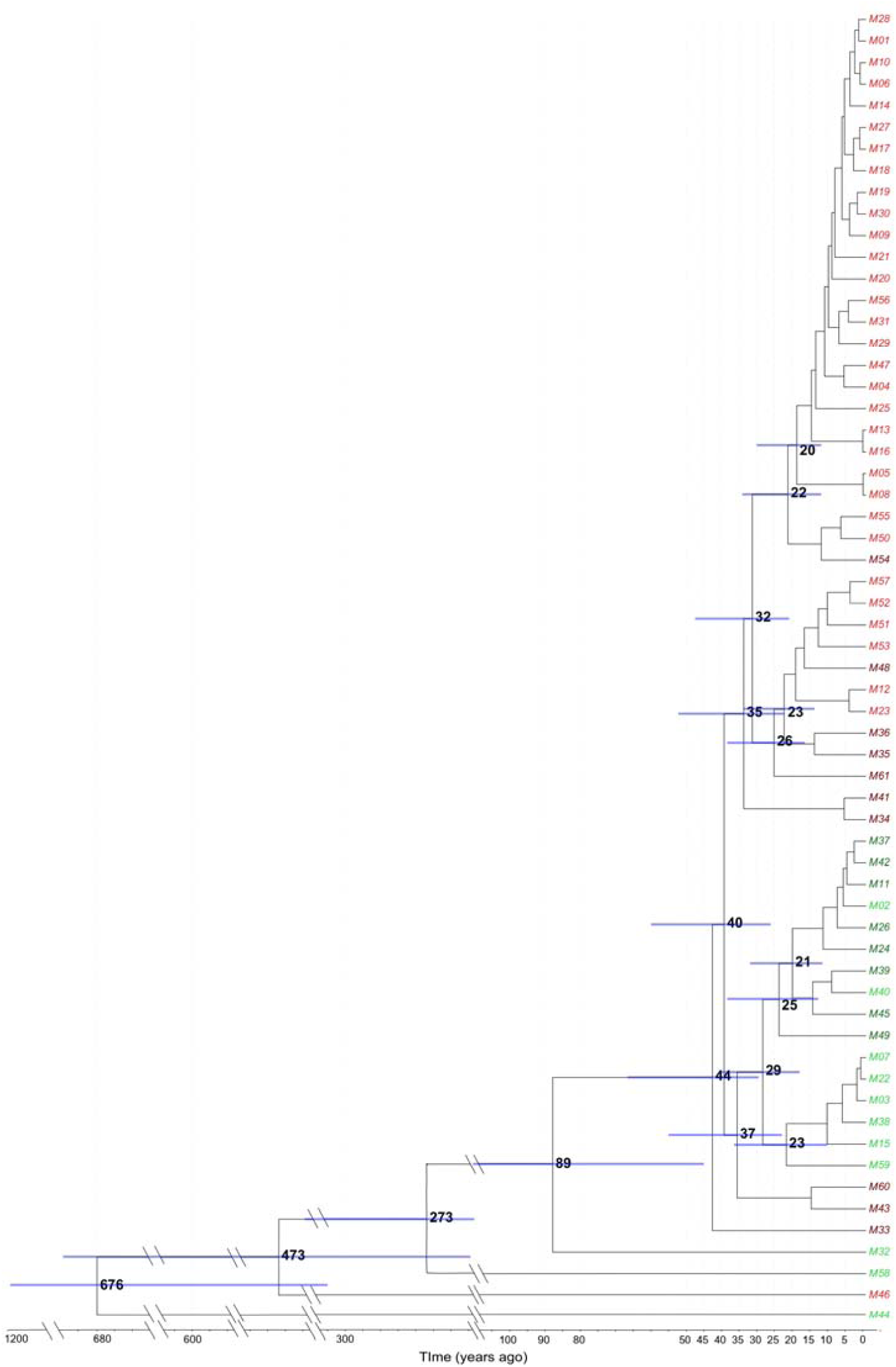
Bayesian maximum clade credibility phylogenetic tree created via BEAST (using single nucleotide polymorphisms) of 61 strains with nodes in increasing order of age. Node ages < 15 years are not shown for clarity. Bars represent 95% HPD interval for node age. Color of tip represents drug susceptibility - Green: pan-susceptible, Dark Red: Resistant only to Isoniazid or Rifampicin, Dark Green: Resistant to a drug other than Isoniazid or Rifampicin, Red: multi-drug resistant.

### Epidemiological vs. genotypic data

We compared the use of genetic and epidemiological data for transmission inference. PE/PPE region differences and indels between closely related pairs were visualized in Integrated Genome Viewer and confirmed to be high confidence (Supplementary methods). Sequencing data was available for three of seven household contact pairs in the cluster of 148 patients. Only one household link (index/parent M23 – contact/child M12) was consistent with a recent transmission event on the tree and by genetic distance (SNP difference = 1). When high-confidence PE/PPE regions were included, the genetic distance between this pair increased to 3 variants. With the further addition of high confidence indels the pairwise distance increased to 4, in the predicted 5 year interval (Supplementary Data 2). No variation was observed in this pair in DR-related loci. The other two isolate pairs from household contacts (index/parent M02 – contact/child M01, index/child M45 – contact/parent M58) were 17 and 421 SNPs apart, respectively. Isolates M01 and M02 were genetically closer to other isolates on the tree (M01-M28 6 SNPs apart and M02-M42-M11-M37 all within 4 SNPs of each other). A pair of isolates collected two months apart from a host who had not been on treatment (MDR strains M06 and M10) was found to have no SNP differences outside of PE/PPE regions. In PE/PPE regions, we found 5 high-quality SNPs; similarly, 2 indels were observed in other regions.

Looking at genetic evidence alone for recent transmission using a distance cutoff of ≤5 SNPs (excluding DR and PE/PPE regions)^3^, 139 links among 38 patients were identified (Figure 3). Other than the pair (index parent M23 and contact child M12), none of these belonged to the same household. With the addition of high confidence indels and PE/PPE SNPs and using the cutoff of ≤7 variants (distance between the two isolates belonging to the same patient when high confidence indels and PE/PPE variants were included), there were 41 links among 30 patients,*i.e*. 76% fewer links than when these variants were excluded. Phylogenetic trees built by including indel variation also had notable differences within the cluster of 57 isolates (Supplementary Figure 3).

**Figure 3:**
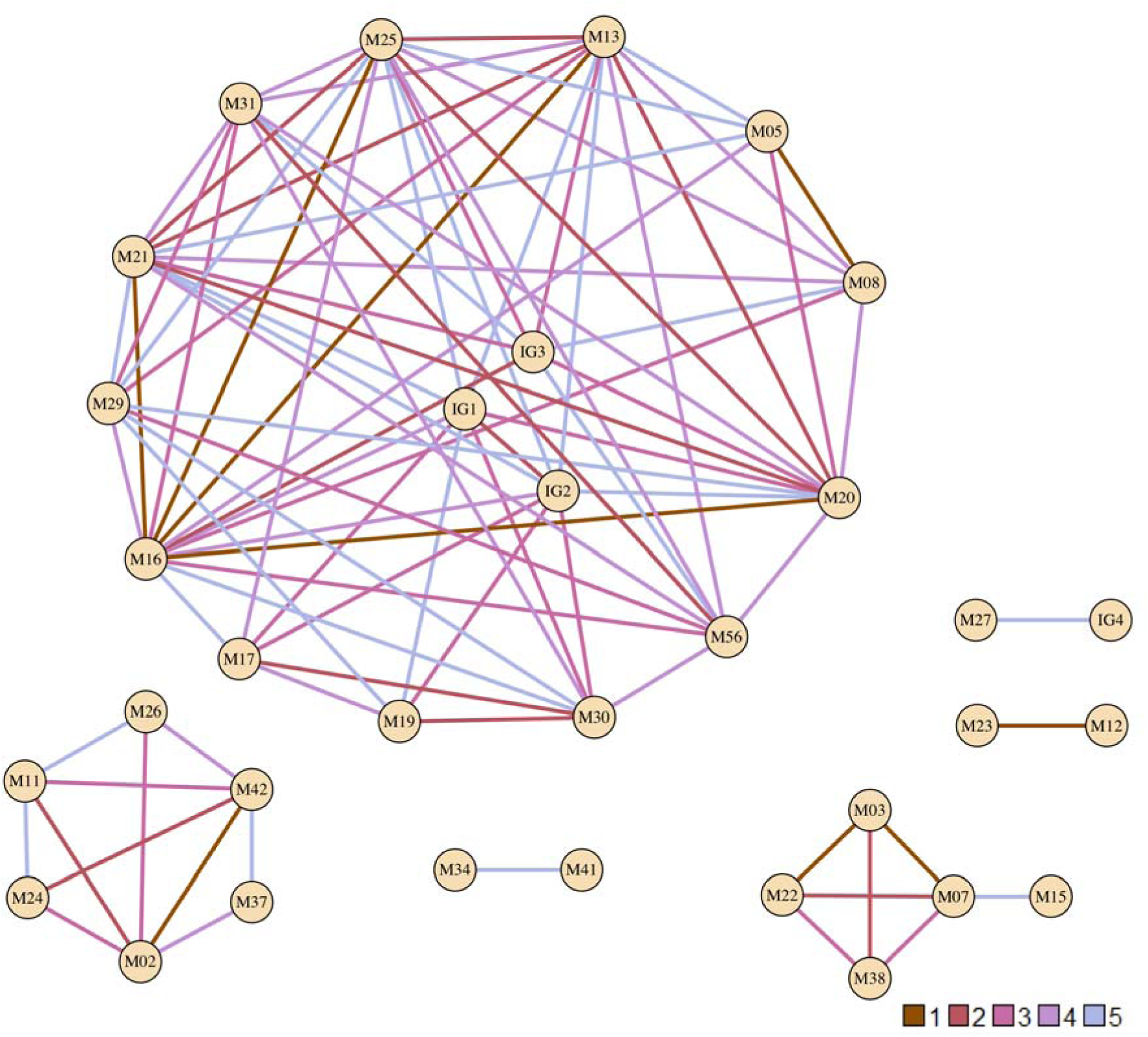
*Network of strains that were less than or equal to 5 single nucleotide polymorphisms (SNPs) apart (with the exclusion of PE/PPE regions and regions coding for drug resistance). IG = Identical Group – consists of strains that had zero SNP difference, IG1 = M09, M14, M01; IG2 = M06, M10; IG3 = M04, M07; IG4 = M18, M28. Legend shows number of different SNPs and corresponding color of the edges. Plot generated using R package igraph*.

Of the 375 isolates sequenced from a TB outbreak in London,^13^ 325 met our quality criteria and were further examined. Using a genetic distance cut-off of ≤5 SNP (excluding DR and PE/PPE regions), 309 of these isolates separated out into one large interconnected cluster consisting of 31,776 links. Among 38 serial isolates that were collected from 19 patients from that outbreak, the largest difference between a pair with the inclusion of indels and PE/PPE regions was also found to be 7 SNPs. When applying this as the threshold for identifying a genetic link, the interconnected cluster was reduced to 294 strains with 28,230 links, i.e. 11% fewer links than when PE/PPE variants and indels were excluded‥

### Host factors

We measured host infectiousness in the cluster using the ‘propensity to propagate’^35^ (PTP) method and identified five patients as having the highest possible score (PTP> 4). This was related to patients being younger (20-29y) males with smear positive PTB and a history of substance use (Supplementary Figure 4). Three of these were identified to be within the network of patients with genetically close MDR isolates (Figure 3). The mean cluster PTP was also high at 1·699.

**Figure 4:**
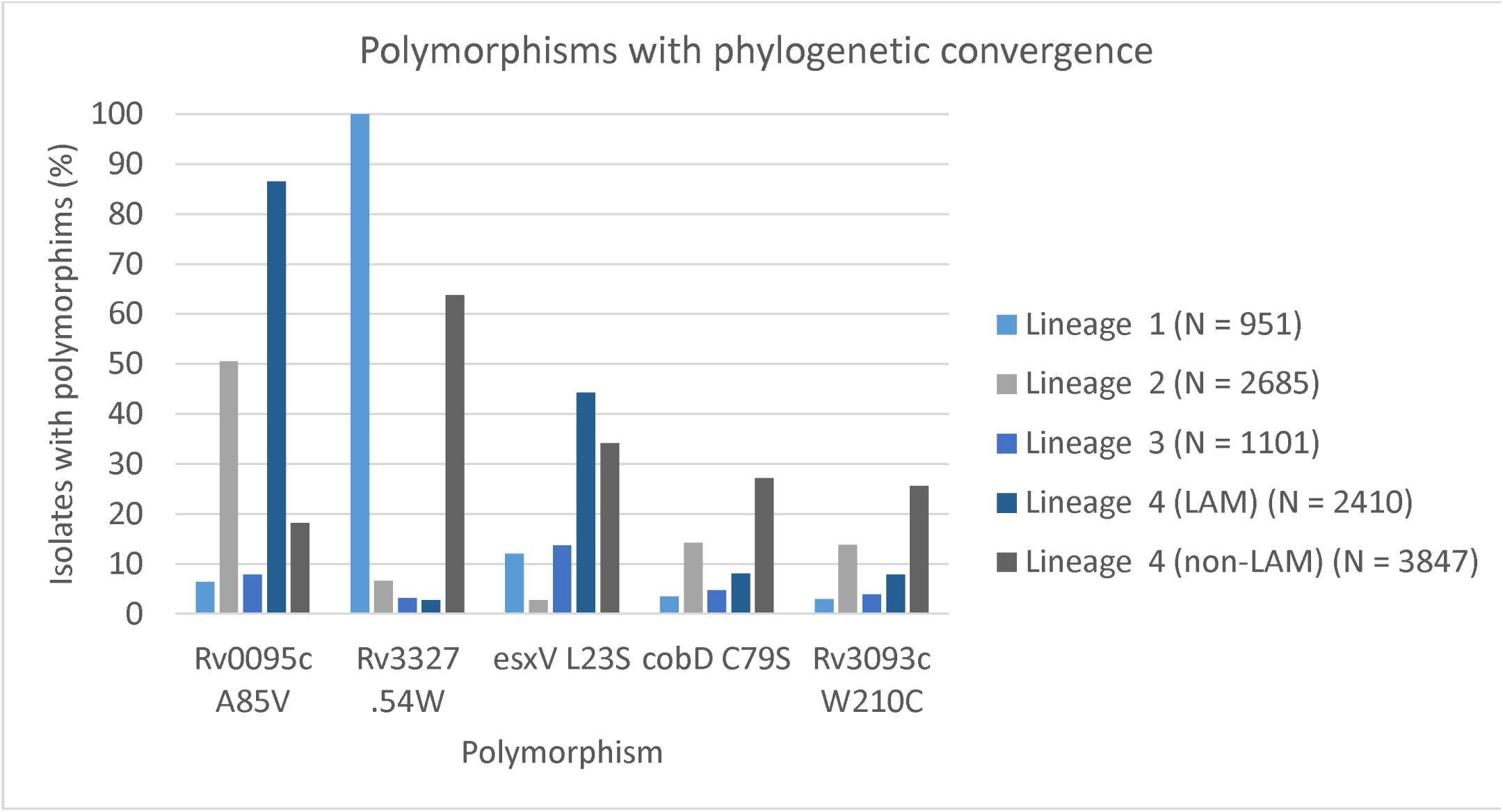
*Single nucleotide polymorphisms in high-transmission cluster of 57 strains showing phylogenetic convergence and their percent frequency among Mycobacterium tuberculosis lineages*.

### Drug Resistance

First and second line drug resistance was acquired several times within the core cluster of 57 isolates (Table 2, Supplementary Figure 5, and Supplementary Table 1).

**Table 2:** Drug resistance acquisition in MTB strains based on phenotype

| **Antibiotic** | **Minimum DR acquisition instances** | **Most recent DR acquisition (year)** |
| --- | --- | --- |
| Isoniazid | 2 | 1972 |
| Rifampicin | 2 | 2007 |
| Pyrazinamide | 5 | 2011 |
| Ethambutol | 3 | 1996 |
| Ciprofloxacin | 1 | 1996 |
| Kanamycin | 1 | 2009 |
| Capreomycin | 1 | 2003 |

*First line drugs:* Isoniazid (INH) resistance was acquired at least twice, and in both times with a *katG* S315T mutation, ca 1968 (95% HPD 1945-1985) and ca 1972 (95% HPD 1952-1985). Rifampicin resistance (RR) was acquired within the large INH resistant clade at least twice ce 1980 (95% HPD 1964-1990, *rpoB* D435V) and ce1986 (95% HPD 1973-1996, most frequent variant *rpoB* S450L). Pyrazinamide (PZA) resistance followed RR and was acquired at least 4 times, the most frequent mutation was *pncA* Q10R acquired ce 1980 (95% HPD 1964-1990), the most recent PZA resistance acquisition event was predicted to be within the last year of isolation (95% HPD 2006-2012). Ethambutol resistance was acquired at least twice within the MDR clade and contemporaneous with RR acquisition in both cases. The mutation Y319S was the most common *embB* mutation observed.

*Second line drugs:* Of the 25 strains tested for ciprofloxacin, one MDR isolate (M43) acquired resistance ce 1996 (95% HPD 1979-2009). Similarly only one (M30, also resistant to capreomycin) of the 41 isolates tested for kanamycin acquired resistance ce 2009 (95% HPD 2006-2012). We were not able to identify a mutations in *gyr* or *rrs* to explain resistance to these two drugs. For capreomycin, seven of forty tested isolates were resistant and carried the *tlyA* G232D mutation estimated to have been acquired ce 2005 (95% HPD 2001-2008).

### Resistance and Bacterial Fitness

As there were several isolates measured to be resistant by the culture-based method that did not harbor any known resistance mutations, e.g. for EMB and PZA, we attempted a phylogeny-based genome wide association within the group of 61 isolates to identify new mutations associated with resistance (Supplementary Table 2). In addition to identifying the known mutations that confer resistance to isoniazid and rifampicin, we found an association between EMB resistance and a mutation (3778221AG) in the intergenic region between *spoU* and *PE-PGRS51* genes (20 bp from *spoU* end and 347 bp before *PE-PGRS51* start), corresponding to the acquisition of EMB resistance 32 years ago shown in Supplementary Figure 5.

We identified 175 mutations that were unique to the core cluster of 57 isolates and were absent from the 4 more distantly related isolates. Hypothesizing that a subset of these mutations may have contributed to transmissibility of this cluster, we measured which are under positive selection by looking at 12,032 other MTB isolates. Five mutations met our criteria for positive selection i.e. were found to have a frequency of >5% in at least 3 other TB lineages (lineage 1,2, 3, and non-LAM-4) (Figure 4). Of the five, two occurred in genes with known function esxV which is an ESAT-6 like secreted protein and cobD, a cobalamin biosynthesis protein.

## Discussion

In this detailed analysis of a MIRU-VNTR cluster with variable degrees of drug resistance from a high prevalence setting, we show that traditional genotyping methods have a significantly lower resolution in identifying transmission clusters as compared to WGS, particularly when variation in PE/PPE regions, indels are incorporated into the analysis. Additionally, we found complex evolutionary patterns within an otherwise identical MIRU cluster and identified the interplay of host and epidemiological factors contributing to transmission potential of a cluster. The higher resolution gained by WGS is consistent with prior reports^2,3,36–38^, but the maximum genetic distance we find between the clustered isolates is larger than previously seen and we further estimate the group of LAM-4.3.3 sequenced isolates to have been circulating for over 8 decades in our study community. Previous studies performing WGS of MIRU-VNTR clusters in low prevalence settings have noted shorter genetic distance between isolates,^2,3,13^ and in one case the distances were insufficient to reliably and consistently inform contact tracing interventions.^13^ It is possible, that certain features of our selected cluster have led to the observation of such high levels of diversity. First our cluster spans the spectrum of pan-susceptible to resistant to 7 drugs, second is that our isolates all belong to lineage 4, a lineage that has been noted to be the most phenotypically and genotypically diverse of the TB lineages.^39^ However, the proportion of diversity that could be linked directly to drug resistance was low. A parsimonious explanation of the high degree of observed genomic diversity is that the rate of MIRU-VNTR pattern evolution is on average slow and on the order of decades. Despite this, MIRU-VNTR likely offers sufficient resolution in low prevalence settings as most TB cases there tend to be imported.^2,3^

The genes in the PE and PPE families constitute about 10% of the TB genome and have been grouped together based on the proline-glutamate (PE) and proline-proline-glutamate (PPE) signature motifs but members of this family are scattered throughout the genome and have diverse functions.^40,41^ Because they carry a high GC content and contain repetitive areas they have been typically excluded from analysis of sequencing data.^42^ However, recent advances in sequencing technology allow for longer read lengths and increased throughput which combined with more accurate bioinformatics pipelines, it is now possible to call variants in a proportion of these genes with high confidence. We identified several high quality indels and variation in the PE/PPE regions in our dataset. The commonly used cutoff of ≤5 SNPs to infer transmission does not take into account the different evolutionary rate of these regions. In our study, when identifying closely related strains with the inclusion of high confidence PE/PPE regions, the number of possible links between strains decreased. An accepted standard that accounts for variation in these regions would allow for improved resolution of transmission events. Although, comparison of ancestral relationships in the phylogenetic trees without and with inclusion of indels did not show significant differences, there were notable differences within the closely related cluster highlighting that similarity measures that rely on SNPs alone could be misleading. Inclusion of indels and PE/PPE regions in estimation of divergence dates is limited by our current lack of knowledge regarding their evolutionary rates but these regions account for an appreciable proportion of variation seen between closely related isolates and thus including this information may affect interpretation of transmission links.^43^

We identified many genomic links using the SNP distance threshold of ≤5 criterion^3^ that were not discovered within household contact investigation, providing evidence that household contact investigation is not sufficient to identify and treat secondary TB cases as transmission can occur anywhere in the community. Additionally, 2 of 3 case pairs that belonged to the same household were found to have large genetic distances making it more likely that transmission occurred outside the household. Although the dataset used was relatively small, these findings add to the current limited literature on the topic.^44–46^ Overall our study highlights the utility of WGS in resolving transmission links particularly in high burden settings where several transmission chains may occur simultaneously. WGS of a well characterized cluster through MIRU-VNTR led to identification of several sub-clusters with further granularity achieved from addition of variants in regions that are routinely excluded from these analyses. With decreasing cost of WGS, sequencing data could be integrated with epidemiological investigation in lieu of traditional fingerprinting methods to identify transmission clusters and for reconstruction of contact networks, particularly given the increasing emphasis on active case finding for TB elimination.^17^

Our phylogenetic dating procedures supports that acquisition of MDR is not recent in Lima, and that MDR cases, given the observed phylogenic structure, are mostly related to transmission. This finding is consistent with other studies performed in other countries including South Africa.^18^ Within the MDR sub-cluster, ethambutol and pyrazinamide was acquired and transmitted. This finding was similar to that of a study in Uganda,^47^ where pyrazinamide resistance typically arose in MDR strains with several different causative mutations. This highlights the importance of testing for pyrazinamide resistance in order to determine benefit of its use in MDR-TB treatment regimens.^48^

Phenotypic resistance could not be explained by genotype in a few isolates including in our study. To this effect, we undertook a GWAS procedure to identify drug resistant phenotype-genotype associations. We identified the intergenic SNP 3778221AG 30bp downstream of the porbable tRNA/rRNA methylase gene *spoU* to be significantly associated with ethambutol resistance. Although a causative mechanism for how this variant modulates EMB susceptibility or fitness is not clear, this finding is supported further by a recent large genome-wide association study of 1452 MTB isolates.^49^

The cluster under study was the largest such cluster observed in the Lima household transmission study. Its transmission success was likely due to both bacterial and host factors. We quantified the host predilection to transmit TB with the PTP measure and found the cluster to have a higher score than the median PTP measure reported by a study in Netherlands.^35^ Our study had five patients with a particularly high PTP above the highest reported value of 3·9,^35^ potentially contributing to transmission in the population we studied. A few prior studies have characterized bacterial genetic factors that contributed to increased transmissibility.^24,27,50^ We add to this literature by identifying five cluster defining SNPs to be under positive selection in a large TB genomic dataset. One of these SNPs (*esxV* S23L) is a member of the ESAT-6 family of secreted proteins, some of which have been shown to be involved in host-pathogen interactions and may thus have contributed to increased transmissibility.^50,51^

Our study had several limitations. Contact tracing was done within household contacts and hence epidemiological links in the community were possibly missed. Sequencing a subset of isolates from the cluster may have led to missed links along the transmission chain. It may also have led to an underestimation of the diversity. However, the sampled subset demonstrated a substantial amount of diversity, more than would be expected within a cluster with identical MIRU pattern.^2,3^ We also cannot exclude that the 2 outer most isolates were miss-assigned the reported MIRU pattern and because of this we focused on the isolates confirmed to be of the same lineage by in silico spoligotyping and the WGS SNP barcode. Finally, it is important to note that our dating estimates are heavily reliant on the molecular clock rate that has been previously reported in the literature.

In summary, our findings add to the evidence challenging the traditional interpretation of a MIRU-VNTR cluster as indicating recent transmission and suggest that the benefits of WGS over MIRU-VNTR may be even more prominent in high prevalence settings when TB transmission has been ongoing without interruption, especially when high confidence PE/PPE and indel genetic variants are included. WGS can also provide insights into biology of MTB to improve our understanding of DR, transmission and host-pathogen interaction.

## Authors’ contributions

Avika Dixit conducted the data analysis, drafted and revised the manuscript.

All authors provided key edits to the manuscript

Additionally:

Luca Freschi contributed to the data analysis

Roger Vargas contributed to the data analysis.

Roger Calderon did the DNA extraction and performed the DST testing

James Sacchettini conducting the sequencing of isolates included in the study

Francis Drobniewski shared genomic and metadata from the London TB outbreak

Jerome T. Galea helped conduct the household contact study in Peru.

Carmen Contreras helped conduct the household contact study in Peru

Rosa Yataco helped conduct the household contact study in Peru.

Zibiao Zhang managed the household contact study Peru data.

Leonid Lecca helped conduct the household contact study in Peru

Sergios-Orestis Koloktronis verified the phylogenetic analysis.

Barun Mathema contributed to study design and phylogenetic analysis.

Maha Farhat conceptualized the study, supervised the data analysis, reviewed and edited the manuscript.

## Declaration of interests

Avika Dixit: None

Luca Freschi: None

Roger Vargas: None

Roger Calderon: None

James Sacchettini: None

Francis Drobniewski: None

Jerome T. Galea: None

Carmen Contreras: None

Rosa Yataco: None

Zibiao Zhang: None

Leonid Lecca: None

Sergios-Orestis Kolokotronis: None

Barun Mathema: None

Maha R. Farhat: None

## Acknowledgments

We thank Megan Murray, the co-PI of the Peru Epi study for helpful input on the manuscript. We thank the patients and their families who contributed to this study. We also thank the Partners in Health healthcare personnel at participating health centers in Lima, Peru. We wish to thank Dr. Robert Husson for his invaluable feedback on an initial draft of the manuscript.

